# Parasitoid community responds indiscriminately to fluctuating spruce budworm and other caterpillars on balsam fir

**DOI:** 10.1101/615799

**Authors:** Christopher J. Greyson-Gaito, Kevin S. McCann, Jochen Fründ, Christopher J. Lucarotti, M. Alex Smith, Eldon S. Eveleigh

**Author notes:** Corresponding Author: (CJGG). **Note:** This version is a preprint and has not been accepted by a peer review process.

## Abstract

The world is astoundingly variable, and individuals to whole communities must respond to variability to survive. One example of nature’s variability is the massive fluctuations in spruce budworm (*Choristoneura fumiferana* Clemens, Lepidoptera: Tortricidae) populations that occur over 35 years. We examined how the parasitoid community altered its parasitism of budworm and other caterpillar species in response to these fluctuations. Budworm and other caterpillar species were sampled from balsam fir in three plots for 14 years in Atlantic Canada, and then reared to identify any emerging parasitoids. We found that the parasitoid community showed a simple linear, indiscriminate response (i.e., no preference, where densities purely dictated parasitism rates) to changes in budworm densities relative to other caterpillar species on balsam fir. We also observed strong changes in topology and distributions of interaction strengths between the parasitoids, budworm and other caterpillar species as budworm densities fluctuated. Our study contributes to the suggestion that hardwood trees are a critical part of the budworm-parasitoid food web, where parasitoids attack other caterpillar species on hardwood trees when budworm populations are low. Taken together, our study shows that a parasitoid community collectively alters species interactions in response to variable budworm densities, fundamentally shifting food web pathways.

## Introduction

Ecologists have long used equilibrium assumptions to study the complex suite of interactions that make up food webs (May 1973, Allesina and Tang 2012). Although a reasonable first approach, in fact, both abiotic and biotic conditions are notoriously variable (Levin 1998, Guichard and Gouhier 2014). However, our understanding of how organisms respond to variation remains surprisingly limited. In light of human-driven impacts including climate change that promise to significantly alter this variation (Cotton 2003, Ims et al. 2008), it behooves ecologists to embrace natural variation and to ask how individuals to whole communities respond to both natural variation and changes in this natural variation caused by human modifications.

Individual and species-level responses to variability can add together to produce community-level and food web responses. Individual and species-level responses can include population changes, colonization/extinction events and behaviour (Supp and Ernest 2014). Indeed, recent work has found compelling evidence that individuals and species behave to take advantage of strongly changing conditions. A fascinating example of individuals responding to changing conditions is grizzly bears in Alaska following the ephemeral pulses of salmon densities across the river and stream landscape (Armstrong et al. 2016). The bears track the phenological variation of salmon spawning across space and time, so maximising the bear’s energy intake. Community-level responses can include richness, evenness, and total biomass changes (Keitt 2008, Supp and Ernest 2014). For example, Supp and Ernest (2014) found that species richness and evenness stayed relatively constant in contrast to species populations when communities were exposed to a disturbance. The intersection of individual and species responses with community responses is an exciting research avenue especially when community and food web responses can be large enough to impact the population dynamics of connected species or ecosystem function.

One known example of a community-level response to variability is the impressive convergence of parasitoids on the periodic spruce budworm (*Choristoneura fumiferana* Clemens, Lepidoptera: Tortricidae) outbreaks on balsam fir (*Abies balsamea* Miller, Pinaceae) (Eveleigh et al. 2007). Budworm have massive and relatively predictable outbreaks every thirty five years, followed by periods of budworm rarity (Royama et al. 2005). This cycle is considered to be a predator – prey cycle, where the predator is a complex of natural enemies including insects that parasitize and then kill a caterpillar host (parasitoids) (Pureswaran et al. 2016, Royama et al. 2017). The diversity of these parasitoids sampled from balsam fir increases when spruce budworm densities increase, similar to how many species of birds converge on a full birdfeeder (the birdfeeder effect) (Eveleigh et al. 2007). The individual parasitoids likely all respond to the changing densities of budworm in order to maximize their fitness (Abrams and Kawecki 1999) and collectively they converge on high densities of budworm, leading to a diversity cascade. The parasitoid community causes between 30-90% mortality depending on the surrounding forest composition and the point in the budworm cycle (Dowden et al. 1950, Cappuccino et al. 1998, Seehausen et al. 2014, Royama et al. 2017). Because the parasitoid community has such a strong response to changing budworm populations and causes high budworm mortality, the budworm – parasitoid food web provides an excellent system to examine community responses to variability.

What is largely unknown about this budworm – parasitoid food web is how the parasitoid community interacts with other caterpillar species in relation to the fluctuations of budworm. When budworm are rare, parasitoid populations decrease, but a reserve population of parasitoids could be maintained by attacking other caterpillar species. Parasitoids, that are predominantly known to consume budworm, switching to attacking other caterpillar species is a possibility because we know some parasitoids are generalists that can successfully attack multiple species (Krombein et al. 1979, Eveleigh et al. 2007, Smith et al. 2011). Therefore, the parasitoid community could converge on high budworm densities during outbreaks and then attack other caterpillar species when budworm densities decline. Specifically, we do not know the relative attack rates of the parasitoid community on budworm and other caterpillar species as budworm densities change. Understanding the contribution of other caterpillar species to the parasitoid community dynamics could help to moderate the amplitude and severity of budworm outbreaks.

Whereas Eveleigh et al. (2007) provided a qualitative examination of the entire budworm food web on balsam fir, and Royama et al. (2017) examined the impact of parasitoids on budworm only, in this exploratory study, we aimed to quantify the changing trophic interactions of parasitoids with both budworm and other caterpillar species on balsam fir during four budworm density phases (before budworm densities peak, during the peak, after the peak, and endemic budworm populations). We analyzed rearing data of budworm and other caterpillar species collected from balsam fir branches sampled from 1982 to 1995. During this time period, balsam fir branches were collected from three plots and a representative sample of budworm and all other caterpillar species were placed into feeding vials to collect and identify any parasitoids that emerged. First, we examined whether the whole parasitoid community exhibited host preference by caterpillar density or type (budworm or other caterpillar species). Host preference was established by testing the relationship between the relative abundances of caterpillars and the relative abundances of parasitoid emergences from each caterpillar type. Second, we evaluated whether the presence or absence of host preference at the level of the parasitoid community was an aggregate response and not a single species response by excluding the most abundant species from calculations of the above relationship. We also examined temporal species diversity turnover to test for an aggregate response. Third, because parasitism rates and species turnover all impact the structure and dynamics of food webs, we examined how the topology and interaction strengths of the budworm food web on balsam fir changed where the number of emergences per caterpillar species was used as interaction strengths. We found that the parasitoid community indiscriminately tracked changes in relative densities of budworm and other caterpillar species on balsam fir, exhibiting a collective response akin to a generalist consumer.

## Materials and methods

### Study sites

Three plots in balsam fir forests in New Brunswick, Canada were established. Plot 1 was in the Acadia Research Forest near Fredericton (46°00’N, 66°25’W). Balsam fir branches were sampled in this plot from 1982 to 1989. Because budworm caused 60% tree mortality in Plot 1 by the mid-1980s, Plot 2 was added, which was also in the Acadia Research Forest. Balsam fir branches were sampled in this plot from 1986 to 1995. In the late 1980s, the budworm populations in Plot 1 and 2 were so low that Plot 3 was added, approximately 170km farther north near Saint-Quentin (47°29’N, 67°15’W). Balsam fir branches were sampled in Plot 3 from 1988 until 1994 when budworm populations also declined to a low level. All plots had mostly balsam fir but also contained spruces and a variety of hardwood trees (Eveleigh et al. 2007). Both the Acadia Research Forest and the Restigouche River watershed (where Plot 3 is located) contained balsam fir dominated, mixed, and hardwood dominated stands (Simard and Clowater 2006, Swift et al. 2006). All plots were outside areas of biopesticide application. Full details of the three plots and all sampling and rearing procedures can be found in Lucarotti et al. (2004), Eveleigh et al. (2007) (SI Materials and Methods) and Royama et al. (2017). Here, we present only a brief synopsis.

### Sampling

At the beginning of each season, a group of codominant balsam fir trees were selected in 20 random locations within each plot. Every year and for each plot, before larval emergence from winter diapause, one balsam fir branch from each of the 20 locations was collected. As soon as second instar larvae in the field began emerging from diapause, balsam fir branches were sampled approximately every day until the end of budworm adult eclosion. On each sampling day during the earlier years when budworm populations were high, one foliated mid-crown balsam fir branch from one of the trees in each of the 20 locations was collected. During the later years when budworm populations were low, two or more branches were collected from each location to increase the number of collected budworm larvae at each sample date and location

### Laboratory work

All budworm and other caterpillar individuals were collected for rearing from all 20 branches sampled before budworm emergence from winter diapause. For branches sampled after budworm emergence from winter diapause, one of the 20 sampled branches was selected and all budworm and other caterpillar individuals on that branch were reared. If a minimum of 100 budworm were obtained for rearing from this branch, no more branches were selected for collection of caterpillars for rearing. If less than 100 budworm were obtained from the first branch selected, then another branch was selected and all budworm and other caterpillar individuals from that branch were collected and reared, even if the final total number of budworm exceeded 100. When budworm populations were low, obtaining more than 100 budworm individuals became difficult. As a result, all budworm and other caterpillar individuals that were found on the sampled branches were collected for rearing. Overall, for every sampling day, all budworm and other caterpillar individuals were reared from a subset of branches of the 20 collected each sampling day. All collected caterpillars (budworm and other caterpillar species) were individually reared on artificial diet (McMorran 1965) and inspected every weekday for mortality. There was high rearing success of both budworm and other caterpillar species because all of these hosts feed on balsam fir and therefore readily feed on the artificial diet. On average, 317 other caterpillar individuals were collected each year. All parasitoids that emerged from any reared caterpillars were morphologically identified to genus and where possible to species. Any parasitoids unidentifiable to at least genus were excluded from our analysis (11% of the total number of emergences from spruce budworm or other caterpillars).

### Statistical Analyses

Because we were interested in quantifying the trophic interactions of parasitoids that attack budworm, we excluded all parasitoid taxa that attacked only other caterpillar species. The 48 parasitoid taxa (listed in Fig. 3) found to attack budworm formed 81% of all recorded trophic interactions with other caterpillar species. Using Chao2 (function specpool, R package vegan, version 2.5.2, (Oksanen et al. 2018)) to estimate the total potential number of interactions between parasitoids and budworm or other caterpillar species, this subsetted dataset captures 74% of the potential interactions between parasitoids and budworm and 63% of the potential interactions between parasitoids and other caterpillar species.

Because budworm populations peaked in different years in the three different plots, we created a new time variable called years before/after peak. In this variable, zero was set as the relative year at which budworm populations peaked in each plot. For all analyses, plots were compared using this relative variable. Hereafter, the phrase **relative year** refers to this created variable “years before/after peak variable”. Plot 1 peaked in 1985 and Plot 3 peaked in 1991. We do not know exactly when budworm populations peaked in plot 2 but because population trends in plots 1 and 2 were nearly identical due to their close proximity, we assumed budworm populations peaked in 1985. We also created a categorical variable called **Peak** with four levels describing the phase of the budworm population cycle each year was in: before the peak (three and two relative years before the peak), during the peak (one relative year before and after the peak, and the peak), after the peak (two and three relative years after the peak), and endemic (four to ten relative years after the peak).

Using the same data, Eveleigh et al. (2007) established through rarefaction that changes in diversity of parasitoid species were not due to sampling artefacts. Consequently, we are confident that any patterns found by the analyses below are not due to changes in sampling intensity but due to underlying ecological mechanisms.

All of the following analyses were done using R version 3.6.3 (R Core Team 2012). The data and the R script used for this manuscript can be found by downloading the Zenodo/GitHub repository at https://doi.org/10.5281/zenodo.1305399 (Greyson-Gaito et al. 2020).

### Parasitoid community host preference

To examine whether the parasitoid community exhibited preference for budworm or other caterpillar species on balsam fir, we calculated two values for every combination of relative year and plot: the ratio of parasitoid emergence from budworm to other caterpillar species for all parasitoid taxa combined, and the ratio of abundances of budworm to other caterpillar species. We ran a generalized least squares (GLS) regression with the log10 of the ratio of emergence as the response variable and the log10 of the ratio of the abundances of budworm to other caterpillar species, plot, and their interaction as the explanatory variables (function gls, R package nlme, version 3.1-145, (Pinheiro et al. 2018)). We fitted the full model using maximum likelihood estimation (ML), and then used backwards selection with likelihood ratio tests (LLRT) to select the final fixed effects. We refitted the final model using restricted maximum likelihood estimation (REML) to give unbiased ML predictors (Zuur et al. 2009). We tested whether the average slope for all plots was significantly different from one and whether the average intercept for all plots was different from zero using one sample t-tests. As per the methods in Greenwood and Elton (1979), a slope different from one indicates density dependent host preference and a intercept different from zero indicates preference for a specific host type (budworm or other caterpillars).

### Aggregate host preference response

To identify whether the presence or absence of host preference at the level of the parasitoid community was driven by a single parasitoid taxon or by the whole community, we examined whether removing abundant parasitoids affected host preference and whether there was turnover in parasitoid taxa over time. To examine whether removing abundant parasitoids affected host preference, first we found the three most frequently emerging parasitoid taxa. Second, we then removed in turn the top parasitoid taxon, the top two parasitoid taxa, and the top three parasitoid taxa from the data. Third, using these three datasets, we ran GLS regressions with the same final model as for the analysis in the parasitoid community host preference analysis. Using one-sample t-tests, we compared the average slopes and intercepts for all plots of each of these models with the average slope and intercepts for all plots produced in the model with all parasitoid taxa included. To examine turnover in parasitoid taxa over time, we ran an nMDS analysis using the Bray-Curtis dissimilarity measure where the abundances of individual taxa were divided by the total number of parasitoid emergences (all taxa) for each relative year and plot (function metaMDS, R package vegan, version 2.5.2, (Oksanen et al. 2018)). We ran a perMANOVA between the four groups in the **Peak** variable (function adonis, R package vegan version 2.5-6). In this perMANOVA, we used the Bray-Curtis dissimilarity measure, constrained permutations within each plot, and maintained the temporal order of permutations.

### Food web topology and interaction strengths

Given the potential for changes in parasitism rates and species turnover to alter food web structure, we examined how topology and interaction strengths changed in the budworm food web on balsam fir. We calculated the number of emergences of each parasitoid taxon from either budworm or other caterpillar species for every relative year. To examine changes in topology, we produced visual bipartite food webs from these numbers of emergences (R package bipartite, version 2.15, (Dormann et al. 2008)). To examine changes in interactions strengths, we calculated the ratio of the median to maximum interaction strengths for every relative year, where the number of emergences was used for interaction strengths. Note, using the number of emergences or the per capita emergences for calculating the ratio of median to maximum interaction strengths yields the same answer. We used the change in ratio of median to maximum interaction strengths to qualitatively assess how the distributions of weak to strong interactions strengths changed over time.

## Results

### Parasitoid community host preference

The final model explaining the log10 ratio of parasitoid emergence from budworm to other caterpillar species included the explanatory variables of the log10 ratio of abundances of budworm to other caterpillars, plot, and their interaction (Log10 budworm to other caterpillar species ratio:Plot interaction, L = 11.429, P = 0.0033, df = 1, log likelihood ratio test, Fig. 1). The average slope for all plots was not significantly different from 1 indicating that the parasitoid community did not prefer either budworm or other caterpillar species in a density dependent manner (0.955*±*0.151, df=15, *P* = 0.771, one-sample t-test). The average intercept for all plots was not significantly different from 0 indicating that the parasitoid community did not prefer either budworm or other caterpillar species in a density independent manner (0.164 *±*0.213, df=15, *P* = 0.454, one-sample t-test).

**Figure 1.**
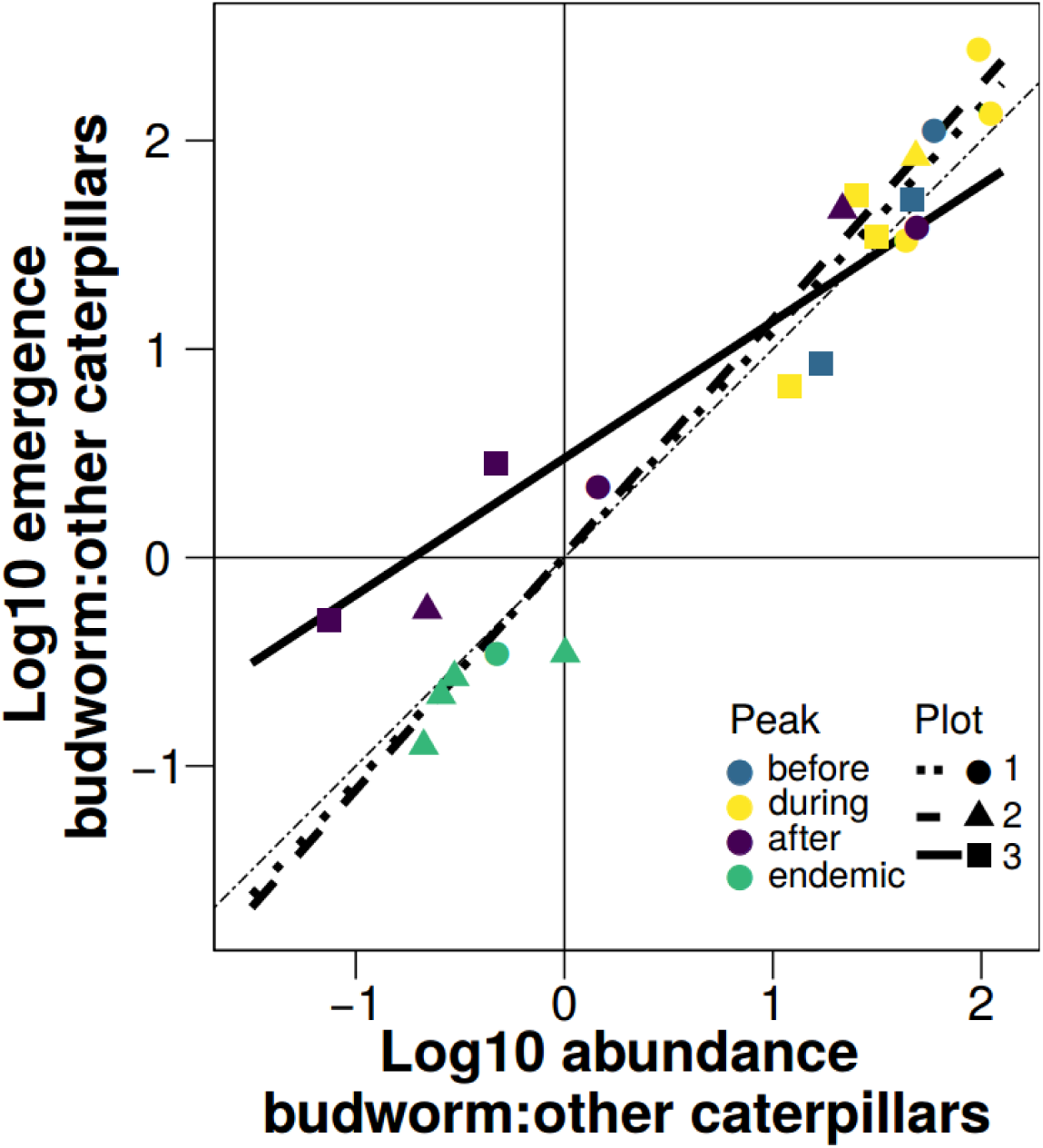
Log10 ratio of parasitoid emergences from budworm to other caterpillar species for all parasitoid taxa used in our analysis as a function of the log10 ratio of all sampled budworm and other caterpillars. Each point is a single relative year and a single plot. The thin dashed line is the y=x line. The parasitoid community did not show a preference for budworm or other caterpillar species by either density or type.

### Aggregate host preference response

Neither dropping the most abundant parasitoid taxon with the most emergences from all caterpillars (*Apanteles fumiferanae* Viereck, Hymenoptera: Braconidae), nor dropping the two most abundant parasitoid taxa (*Apanteles fumiferanae* and *Glypta fumiferanae* Viereck, Hymenoptera: Ichneumonidae), nor dropping the three most abundant taxa (*Apanteles fumiferanae, Glypta fumiferanae*, and *Smidtia fumiferanae* Tothill, Diptera: Tachinidae) caused the average slopes and intercepts for all plots to be significantly different from when all parasitoid taxa were included (original average slope was 0.955 and original average intercept was 0.164, Table 1). However, as each most abundant parasitoid taxon was dropped, there was a trend for decreasing slopes. The parasitoid community did not differ between before and during the peak, but the parasitoid community in these two periods did differ from after the peak and during the endemic periods (F = 5.918, *P* = 0.003, 999 permutations, perMANOVA, Fig. 2).

**Table 1.** Slopes and intercepts with corresponding standard errors, t statistics, p values, and degrees of freedom when the three most abundant parasitoid taxa were dropped consecutively. The explanatory variables in this model were Log10 ratio of abundance of budworm to other caterpillars, plot, and their interaction. The response variable was Log10 ratio of emergences from budworm to other caterpillars.

| Dropped taxa | slope | slope<br>SE | slope<br>t | slope<br>P | intercept | intercept<br>SE | intercept<br>t | intercept<br>P | df |
| --- | --- | --- | --- | --- | --- | --- | --- | --- | --- |
| A. fumiferanae | 0.864 | 0.321 | -0.558 | 0.585 | -0.038 | 0.454 | -0.546 | 0.593 | 15 |
| A. fumiferanae &<br>G. fumiferanae | 0.804 | 0.380 | -0.779 | 0.448 | 0.002 | 0.538 | -0.589 | 0.564 | 15 |
| A. fumiferanae &<br>G. fumiferanae &<br>S. fumiferanae | 0.757 | 0.413 | -0.942 | 0.361 | -0.011 | 0.584 | -0.586 | 0.566 | 15 |

**Figure 2.**
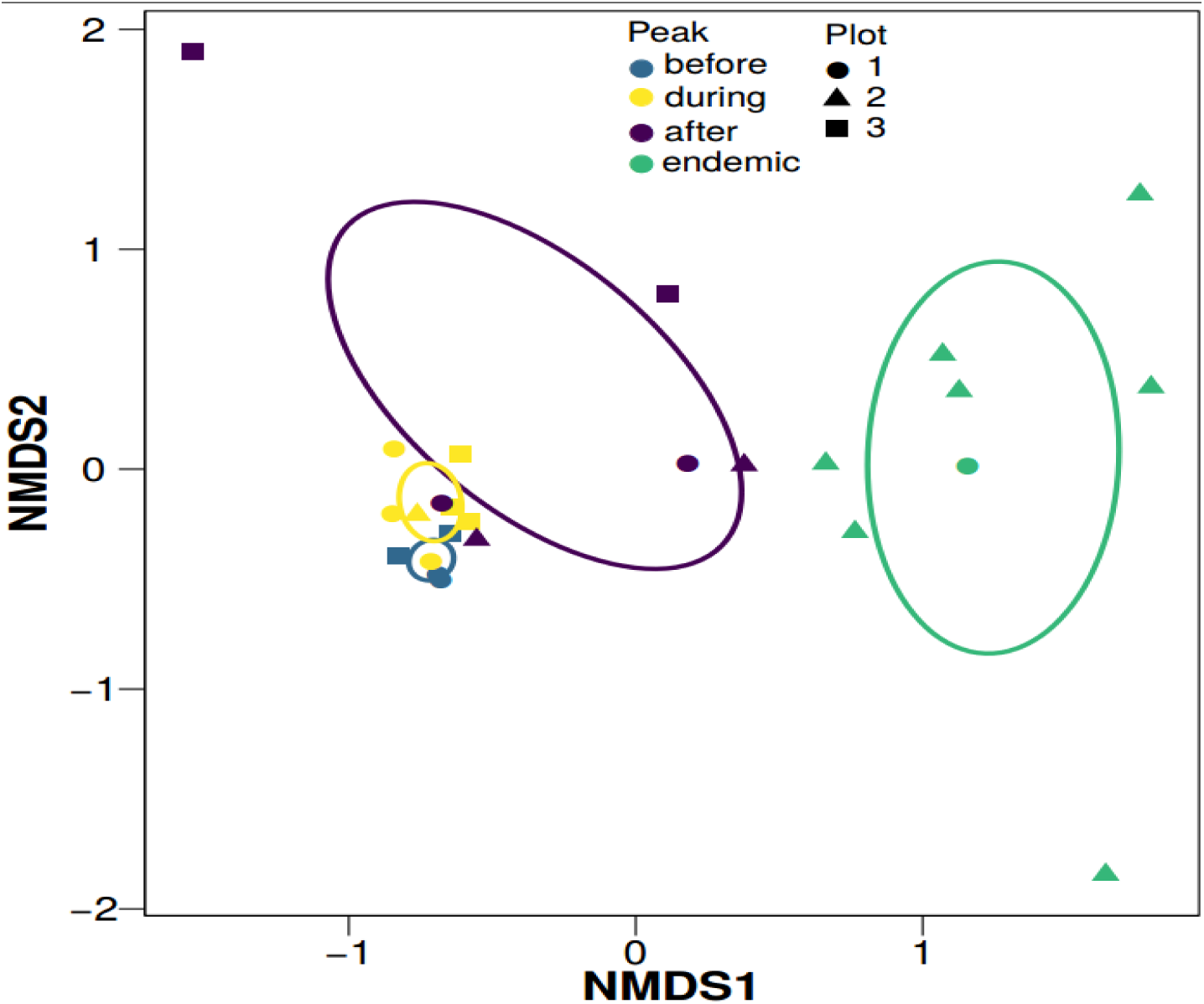
nMDS of parasitoid community emerging from budworm and other caterpillar species on balsam fir over time. The colour of each point and ellipse corresponds to the four temporal groups: three and two relative years before the peak (before – average budworm abundance 7296); one relative year before and after the peak, and the peak (during – average budworm abundance 8067); two and three relative years after the peak (after – average budworm abundance 1128); and four to ten relative years after the peak (endemic – average budworm abundance 29). Each point is a single relative year and a single plot. Each ellipse is a covariance ellipse. 20 iterations. Final stress of 0.087. Instability for preceding 10 iterations was 0.0111. The parasitoid communities before and during the peak were significantly different from after the peak.

**Figure 3.**
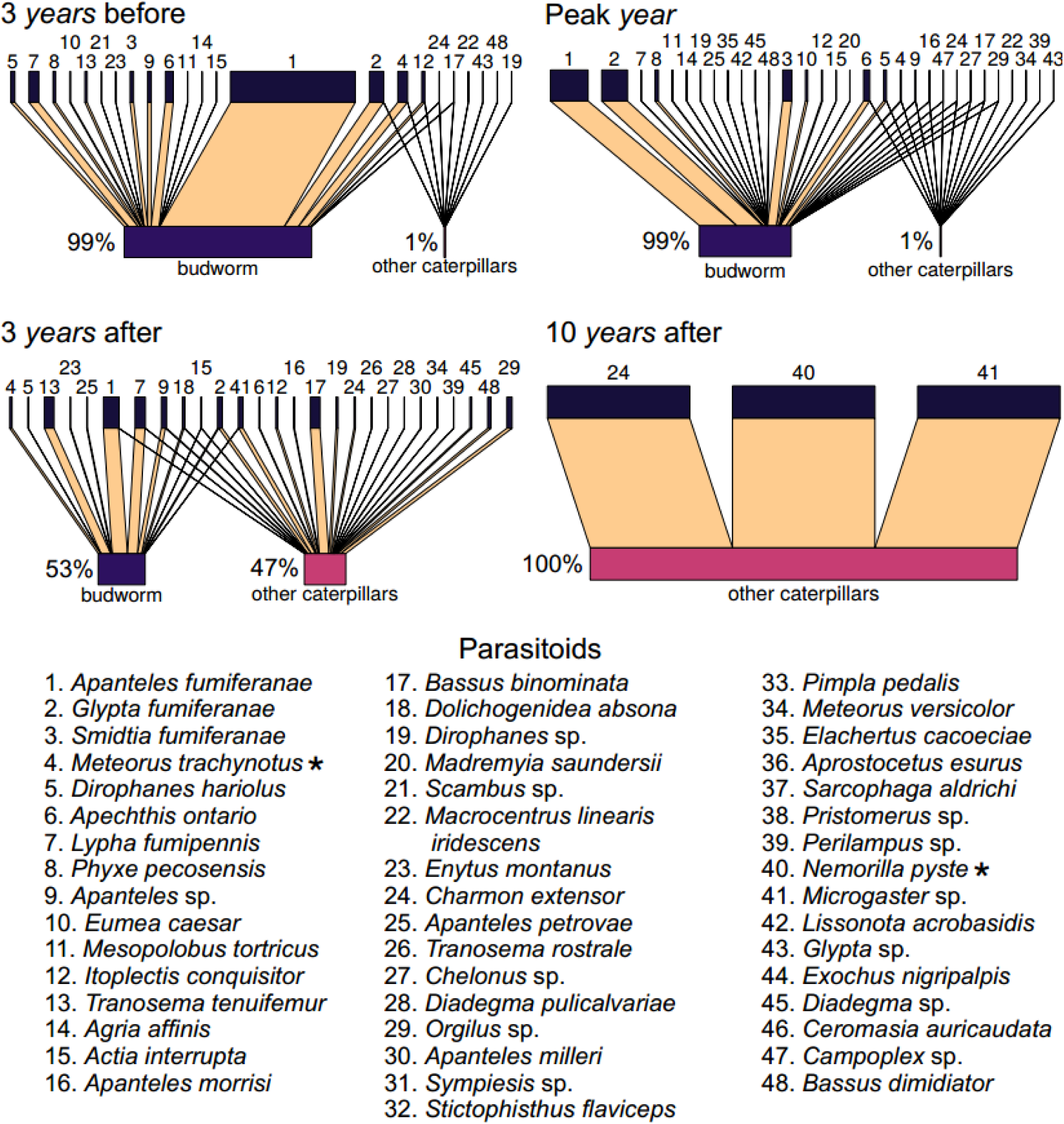
Graphical representations of the number of emergences of each parasitoid taxon (top boxes) from budworm and other caterpillar species (bottom boxes) over time. The width of links is proportional to the fraction of emergences of each parasitoid taxon from either budworm or other caterpillars. The width of the bottom boxes are proportional to the number of emergences from budworm versus other caterpillars, and the percentages show this quantitatively. Four different relative years are shown, where all plots were combined within a relative year: (A) three relative years before the peak, (B) peak relative year, (C) three relative years after the peak, and (D) ten relative years after the peak. All other relative years can be found in Figs S4 & S5. A star denotes a taxon that requires an alternate caterpillar host to overwinter in. To find the corresponding taxon in Eveleigh et al. (2007), see Table S1.

### Food web topology and interaction strengths

There were some parasitoid taxa (e.g. *Diadegma pulicalvariae* Walley, Hymenoptera: Ichneumonidae) that were ephemerally found in the food web (Figs 3, S4, & S5). Parasitoid taxa that were found in the food web consistently through time (e.g. *Apanteles fumiferanae*), often changed from emerging from both budworm and other caterpillar species or just one caterpillar type within a year. The distribution of interactions strengths for budworm and other caterpillar species changed from a skewed distribution dominated by weak interactions towards a uniform distribution, though the variation in the median:maximum interaction strengths between sequential years is greater for other caterpillar species than for budworm (Fig. 4).

**Figure 4.**
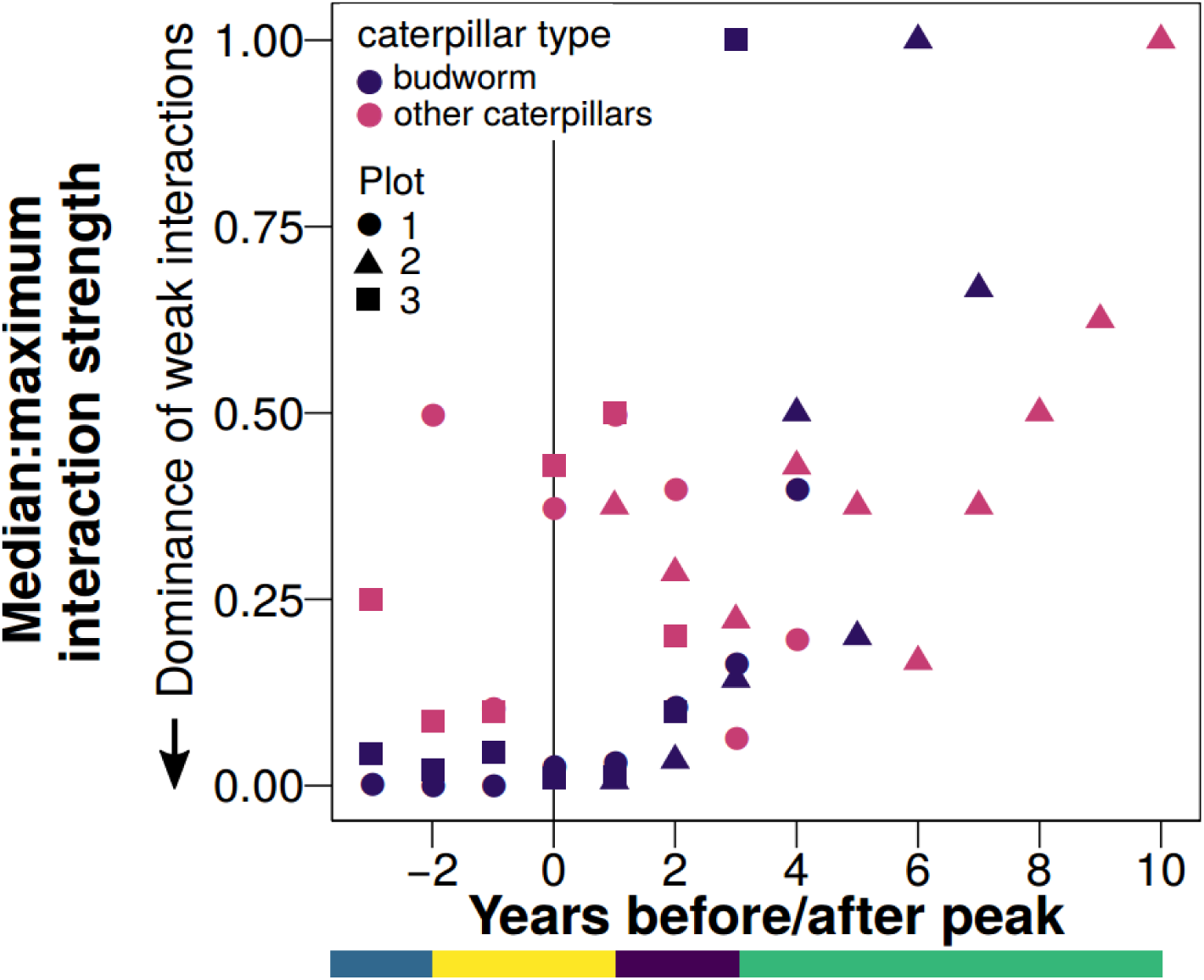
Median:maximum interaction strength over time, for each plot and for each caterpillar type, where the number of emergences was used for interaction strengths. Figure inspired by the median:maximum interaction strength figure in Ushio et al. (2018). Bar at bottom depicts the Peak variable level each year is in: (going from left to right) before, during, after, endemic. As budworm densities decreased, the distribution of interaction strengths shifted from a dichotomy of strong and weak interaction strengths but skewed with a preponderance of weak interactions to a uniform distribution of interaction strengths.

## Discussion

In our study, we have shown that this boreal insect food web is highly responsive and flexible in time to changing budworm densities. We used a 14 year long-term dataset of host/parasitoid abundance and diversity to assess how parasitism rates and trophic interactions changed over the course of a budworm cycle. We found an aggregated whole community correspondence of parasitism rates with caterpillar relative density (budworm:other caterpillar species density) and a sharp change in topology and interaction strength distributions on balsam fir as budworm densities fluctuated.

Interestingly, we found that the parasitoid community overall did not show a preference for budworm or other caterpillar species by either density or type (Fig. 1). This pattern suggests that the parasitoid community indiscriminately attacks budworm and other caterpillar species on balsam fir. Seemingly, the parasitoids do not differentiate between budworm and other caterpillar species populations on balsam fir trees. Instead, they attack whatever is available and parasitoid emergences follow the relative densities of budworm and other caterpillar species. One implication of this parasitoid community indiscriminate response is for modelling of the budworm system. A key aim of theoretical modeling is to simplify the complexity of nature (Yodzis 1989). When the parasitoid community emergences follow the relative densities of budworm and other caterpillars, we can use a single variable for the parasitoid community instead of many variables for each parasitoid species. Furthermore, the consumption function of the parasitoid community on each caterpillar type on balsam fir could be a simple, linear consumptive function. These simplifications seem realistic considering our analysis and would increase the tractability of theoretical modeling of budworm dynamics.

The indiscriminate response by the whole parasitoid community could be caused either by a few dominant parasitoid taxa or be a summation of all parasitoid taxa responses. When we excluded the three most abundant parasitoid taxa from our dataset, the resultant parasitoid communities still largely exhibited no host preference by density or by type, though the slopes did decrease slightly indicating decreased density dependent host preference for budworm. The less common parasitoids seemingly have a greater preference for other caterpillar species than the common parasitoids which is corroborated by examining the preferences of each of the three most common parasitoids; *Apanteles fumiferanae* emerged from budworm more than other caterpillar species regardless of the relative frequencies of budworm and other caterpillar species (Fig. S1), *Glypta fumiferanae* emerged from caterpillars (both budworm and other caterpillars) when budworm were abundant but were generally not found when budworm were rare (Fig. S2), and *Smidtia fumiferanae* emerged from only budworm (Fig. S3). This indicates that parasitoid taxa have differing preferences for budworm and other caterpillars, but collectively, the community exhibits no preference. Differing preferences of each parasitoid taxon could produce species turnover over time and indeed we did find species turnover (Fig. 2). Further support of differing preferences leading to species turnover comes from Royama et al. (2017), who found that no single parasitoid functional group determined the yearly budworm cycle. Instead, as budworm densities changed, there was turnover in the parasitoid functional group that attacked budworm the most, which produced a relatively constant overall parasitism rate of budworm. As a possible mechanism, Royama et al. (2017) posited that the relative profitability of budworm and other caterpillar species changes in time differently for each parasitoid species, where profitability is defined as the relative energy content plus the number of caterpillars that can be attacked for a given amount of hunting effort. Consequently, different parasitoid species would attack budworm at different time periods during the budworm cycle. Theoretical work supports this supposition where two consumers attack a common resource at different rates during the cycling of the resource (Armstrong and McGehee 1980, Xiao and Fussmann 2013). Overall, our results suggest that the parasitoids act individually but produce a unified response to fluctuating budworm densities.

The observed large changes in parasitism rates and species turnover appeared to translate into fluctuating topology and interactions strengths of the food web. We found large changes in topology with many parasitoid taxa emerging from budworm and/or other caterpillar species in some years and not others (Fig. 3). We also found shifts in the distribution of interaction strengths over the budworm cycle (Fig. 4). When budworm were at high densities, the distribution of interaction strengths showed a dichotomy of strong and weak interaction strengths but skewed with a preponderance of weak interactions. As budworm densities declined, the distribution of interaction strengths became uniform. We acknowledge that spatial sampling effort (in terms of number of plots) differs between years and because different plots were sampled at different times along the budworm cycle, plot identity may impact the interactions found. However, in a subset of plots that were sampled in the same relative years (−3 to 4), we still see a trend of increasing median:maximum interaction strength. Therefore, we would argue that even if we had sampled balsam fir in all three plots from budworm population peak to trough, then we would still find a change from skewed interaction strengths distributions to uniform. Similarly, Ushio et al. (2018) found that interaction strengths distributions in a marine fish food web were dominated by weak interactions in the summer and were more uniform in the winter. Ushio et al. (2018) posited that higher productivity in the summer months leading to higher fish abundance drove these fluctuations in interaction strength distributions. Greater budworm densities could be thought of as the same as high fish abundance in the summer. Given the argument for weak interactions as a major stabilising mechanism in a diverse food web (McCann et al. 1998, Gellner and McCann 2016), finding weak interactions dominating during high productivity periods in both the budworm and marine fish food webs is intriguing because these high productiviy periods may be a temporal period that most requires stabilisation (Rosenzweig 1971, Mougi and Nishimura 2007). The other drivers posited by Ushio et al. (2018) include behavioural or physiological responses that vary over time. We suggest too that the behavioural responses by individual parasitoids are integral to the fluctuations in interaction strength distributions found in the budworm food web.

Theory centred on behavioural responses to variable resources may help to explain the observed aggregated simple linear community response and changes in interaction strength distributions. One theoretical model proposes that higher trophic level generalist consumers react to variation in their resources by either increasing consumption of a resource in one separated subgroup of an entire food web (coupling to a resource compartment) or decreasing consumption of a different resource in another separate subgroup of the entire food web (decoupling from a resource compartment) (McCann et al. 2005, McMeans et al. 2016). This coupling and decoupling of different resource compartments can mute large population variation in lower trophic level organisms and so can stabilize food webs. In the budworm – parasitoid food web, although individual parasitoid species may be specialists or generalists, the aggregate response suggests that the collective parasitoid community could be seen as a generalist consumer. The parasitoid community “generalist consumer” couples the resource compartment with balsam fir as the basal resource (hereafter referred to as balsam fir resource compartment) when budworm populations are increasing, and decouples the balsam fir resource compartment when budworm populations are decreasing. The parasitoid community “generalist consumer” seemingly though does not differentiate between budworm and other caterpillar species on balsam fir. Regardless, this hypothesis for the parasitoid community response requires another resource compartment separate from the balsam fir resource compartment.

We suggest that the other resource compartment in the budworm – parasitoid food web has hardwood trees as the basal resource, where white birch (*Betula papyrifera* Marshall, Betulaceae) and red maple (*Acer rubrum* Linnaeus, Sapindaceae) are hardwood trees. Suggestions for this supposition come from previous studies. First, Eveleigh et al. (2007) and Smith et al. (2011) using the same plots as in this study found different responses of parasitoids in plot 3 compared to plots 1 and 2. Eveleigh et al. (2007), using morphological methods, and Smith et al. (2011), using DNA barcoding methods, found greater parasitoid diversity in plot 3 compared to plots 1 and 2. Plot 3 had a lower dominance of balsam fir compared to plots 1 and 2, but plots 2 and 3 had equal proportions of hardwood trees (Eveleigh et al., 2007, SI Table 1). Second, there have been several observations that budworm densities in stands that contained a mixture of softwoods and hardwoods, otherwise known as mixed forest stands, were lower than budworm densities in balsam fir dominated stands during an outbreak, even after accounting for tree densities (Su et al. 1996, Cappuccino et al. 1998, Eveleigh et al. 2007). Consequently, these researchers hypothesized that there must be greater diversity and abundances of parasitoids in mixed forest stands, maintained by the greater diversity and abundances of caterpillar hosts in mixed forest stands over the full duration of a budworm population cycle. This mixed stand hypothesis is related to the hardwood resource compartment hypothesis where both are positing that hardwood trees play a major role for the parasitoid community that attacks spruce budworm by providing alternate and alternative caterpillars. Indeed, our study shows that other caterpillar species are important to the parasitoid community that attacks budworm suggesting that the mixed stand and hardwood resource compartment hypotheses are mechanistically feasible. However, other caterpillar species are chronically undersampled preventing a clear test of these interrelated hypotheses. Even our study undersamples the interactions between other caterpillar species on balsam fir and parasitoids (using Chao2 with interactions instead of species, 63% of the potential interactions between parasitoids and other caterpillar species were sampled in this study). The interactions of other caterpillar species on hardwoods with parasitoids are sampled even less. Consequently, to fully understand budworm dynamics, it is imperative to sample the interactions of parasitoids with other caterpillar species on balsam fir and hardwoods and to ascertain the parasitoid community’s response to other caterpillar species on hardwoods.

The parasitoid community response to changing budworm populations illustrates the fantastic flexibility of food webs. Previous research found that as budworm densities increase on balsam fir, the diversity of parasitoid species found on balsam fir increase at all trophic levels (Eveleigh et al. 2007). In times of budworm rarity, parasitoid species diversity on balsam fir drops and yet the parasitoid community must be maintained by some mechanism otherwise the swift parasitoid community response to increased budworm abundance could not occur. Our study revealed that the parasitoid community responded to changing densities of budworm by linearly and indiscriminately following the relative densities of budworm and other caterpillar species on balsam fir. Large changes in topology and interaction strengths in the budworm food web on balsam fir resulted from the changes in parasitism rates and species turnover. The other caterpillar species that these parasitoids attack are not solely found on balsam fir, and in fact, many researchers have suggested that caterpillars on hardwoods should be the dominant resource while budworm are rare (Su et al. 1996, Cappuccino et al. 1998, Eveleigh et al. 2007). Consequently, further research should include caterpillars on hardwoods and could identify whether the parasitism rates of budworm on balsam fir compared to the parasitism rates of caterpillars on hardwoods change as budworm densities peak and ebb away. Such a response, which appears to be created by the combined actions of all parasitoid species, is an excellent example of community ecology driving the population ecology of a dominant species. For budworm management, we have highlighted the importance of examining whether parasitoids attack other caterpillar species on hardwoods which could mute the amplitude of budworm outbreaks, helping to reduce the defoliation and destruction of balsam fir forests in eastern North America.

## Supporting information

Supporting Information for manuscript (5 figures, 1 table)

## Acknowledgements

We thank the members of the McCann and Eveleigh laboratories for their comments. We are grateful to the many experts for their aid in identifying the insect parasitoids. These experts were J. Barron, A. Bennett, H. Goulet, J. Huber, J. O’Hara, M. Wood, and M. Sharkey. None of this research would have been possible without the many technicians who painstakingly sorted the balsam fir branches searching for caterpillars. Financial support was provided by the Canadian Forest Service to E.S. Eveleigh and C.J. Lucarotti, by the Natural Sciences and Engineering Research Council of Canada to K. S. McCann, and M. A. Smith, by the Atlantic Canada Opportunities Agency to M. A. Smith and Eveleigh, and by the German Research Foundation (DFG, FR 3364/1-1) to J. Fründ

## Author’s Contributions

ESE designed the initial study. ESE and CJL performed the field and laboratory work. CJGG and JF did the statistical analysis with assistance from ESE, MAS, and KSM. CJGG wrote the first draft of the manuscript. All authors contributed to editing the manuscript.

## Data accessibility

The data and the R script used for this manuscript can be found by downloading the Zenodo/ GitHub repository at https://doi.org/10.5281/zenodo.1305399.

