## Supporting Information for manuscript (5 figures, 1 table) for "Parasitoid community responds indiscriminately to fluctuating spruce budworm and other caterpillars on balsam fir"

**ORCID:**
CJGG – 0000-0001-8716-0290
KSM – 0000-0001-6031-7913
JF – 0000-0002-7079-3478
CJL – 0000-0002-3490-568X
MAS – 0000-0002-8650-2575
ESE – 0000-0001-5060-8565

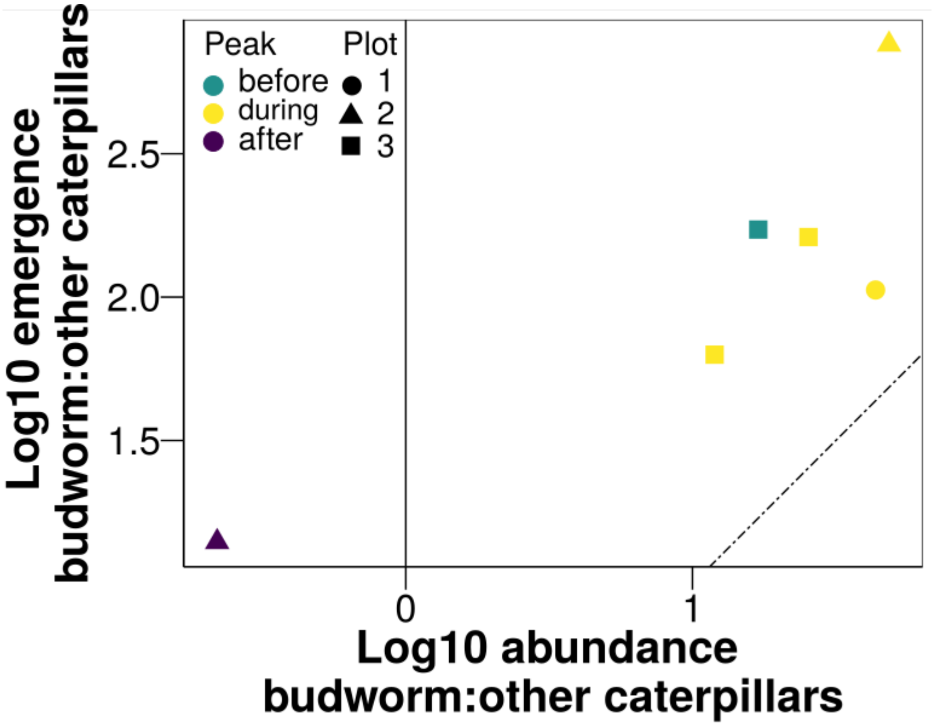


Figure S1 Log10 ratio of *Apanteles fumiferanae* emergences from budworm to other caterpillar species as a function of the log10 ratio of all sampled budworm and other caterpillars. Each point is a single relative year and a single plot. The thin dashed line is the y = x line.

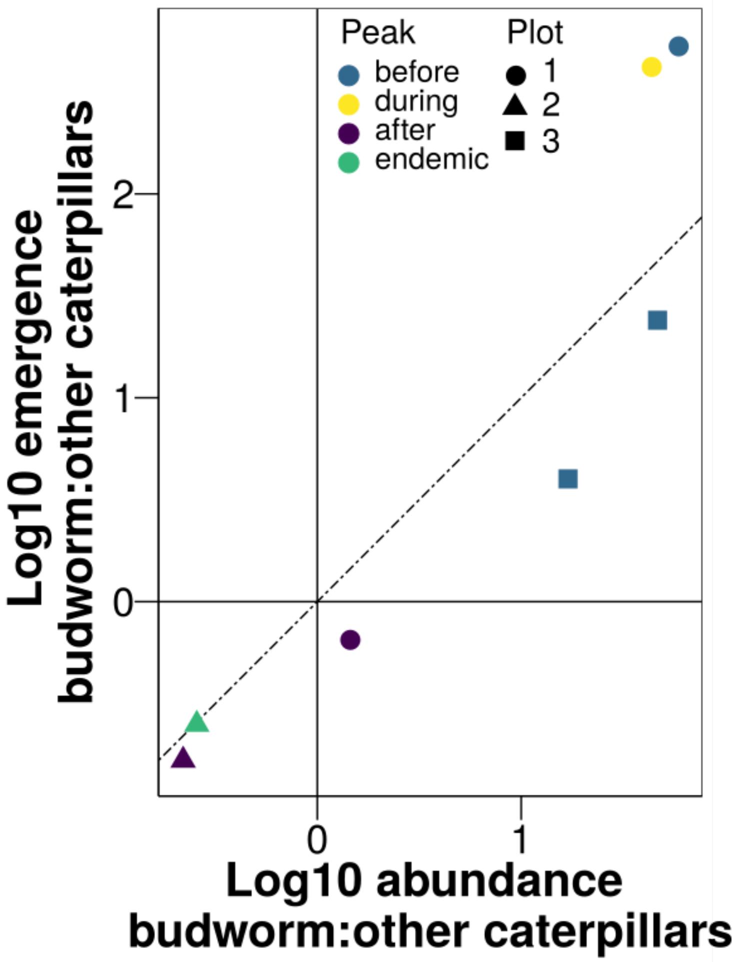


Figure S2 Log10 ratio of *Glypta fumiferanae* emergences from budworm to other caterpillar species as a function of the log10 ratio of all sampled budworm and other caterpillars. Each point is a single relative year and a single plot. The thin dashed line is the y = x line.

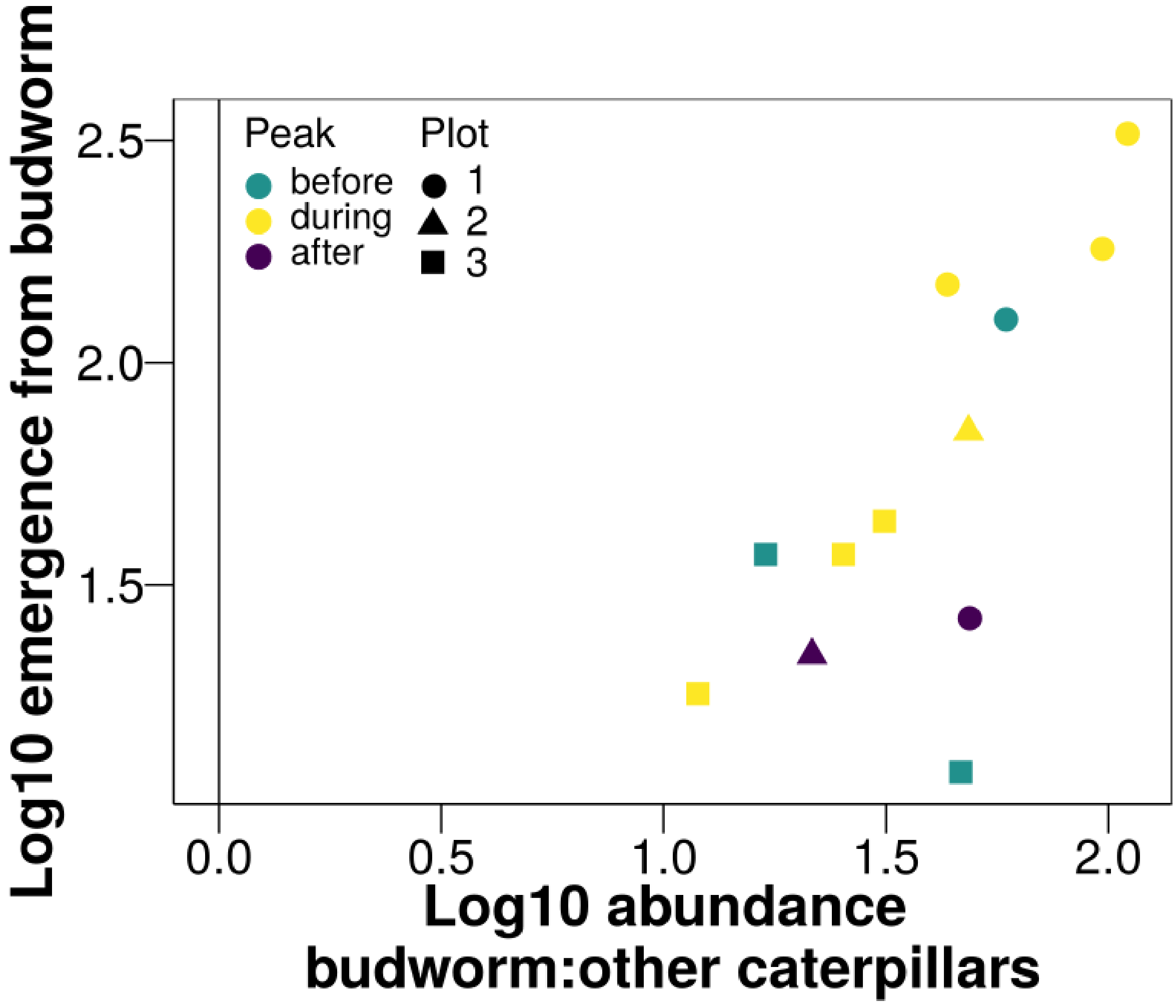


Figure S3 Log10 of *Smidtia fumiferanae* emergences from budworm as a function of the log10 ratio of all sampled budworm and other caterpillars. Each point is a single relative year and a single plot. Note in this dataset, *Smidtia fumiferanae* did not emerge from other caterpillar species and so a ratio of emergences from budworm to other caterpillar species can not be calculated.

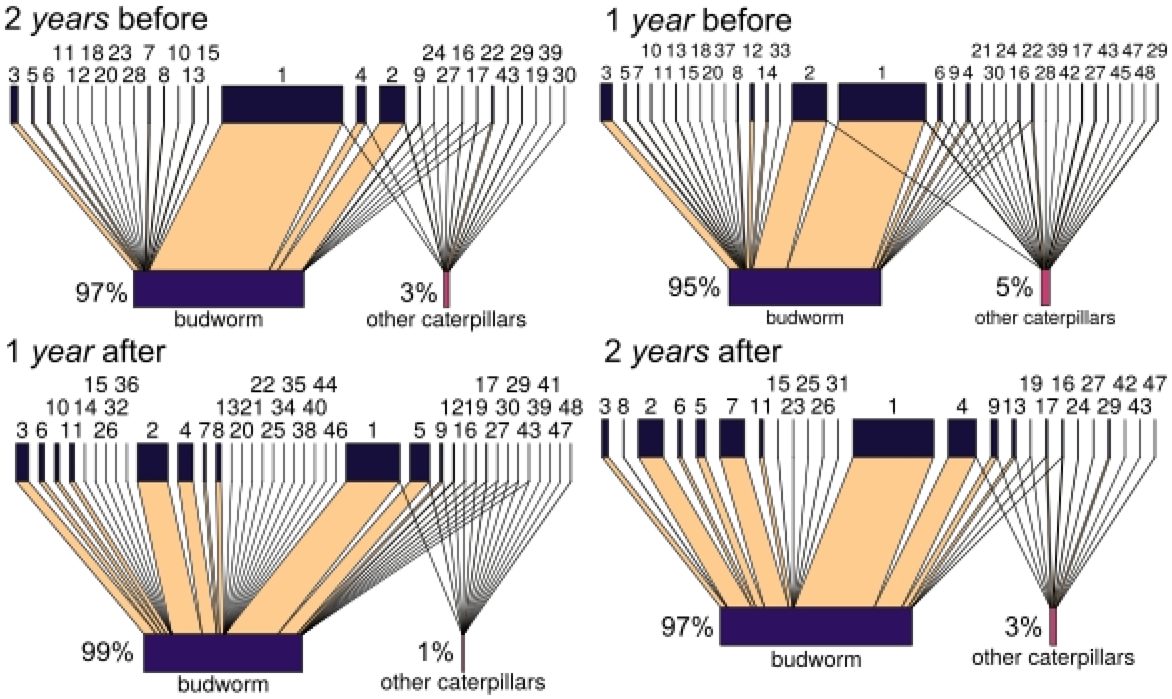


Parasitoids

1. *Apanteles fumiferanae*
2. *Glypta fumiferanae*
3. *Smidtia fumiferanae*
4. *Meteorus trachynotus* ★
5. *Dirophanes hariolus*
6. *Apechthis ontario*
7. *Lypha fumipennis*
8. *Phyxe pecosensis*
9. *Apanteles* sp.
10. *Eumea caesar*
11. *Mesopolobus tortricus*
12. *Itoplectis conquisitor*
13. *Tranosema tenuifemur*
14. *Agria affinis*
15. *Actia interrupta*
16. *Apanteles morrisi*
17. *Bassus binominata*
18. *Dolichogenidea absona*
19. *Dirophanes* sp.
20. *Madremyia saundersii*
21. *Scambus* sp.
22. *Macrocentrus linearis iridescens*
23. *Enytus montanus*
24. *Charmon extensor*
25. *Apanteles petrovae*
26. *Tranosema rostrale*
27. *Chelonus* sp.
28. *Diadegma pulicalvariae*
29. *Orgilus* sp.
30. *Apanteles milleri*
31. *Sympiesis* sp.
32. *Stictophisthus flaviceps*
33. *Pimpla pedalis*
34. *Meteorus versicolor*
35. *Elachertus cacoeciae*
36. *Aprostocetus esurus*
37. *Sarcophaga aldrichi*
38. *Pristomerus* sp.
39. *Perilampus* sp.
40. *Nemorilla pyste* ★
41. *Microgaster* sp.
42. *Lissonota acrobasidis*
43. *Glypta* sp.
44. *Exochus nigripalpis*
45. *Diadegma* sp.
46. *Ceromasia auricaudata*
47. *Campoplex* sp.
48. *Bassus dimidiator*

Figure S4 Graphical representations of the number of emergences of each parasitoid taxon (top boxes) from budworm and other caterpillar species (bottom boxes) over time. The width of links is proportional to the fraction of emergences of each parasitoid taxon from either budworm or other caterpillars. The width of the bottom boxes are proportional to the number of emergences from budworm versus other caterpillars, and the percentages show this quantitatively. Four different relative years are shown, where all plots were combined within a relative year: two relative years before the peak, one relative year before the peak, one relative year after the peak, and two relative years after the peak. A star denotes a species that requires an alternate caterpillar host to overwinter in. To find the corresponding species in Eveleigh et al. (2007), see Table S1.

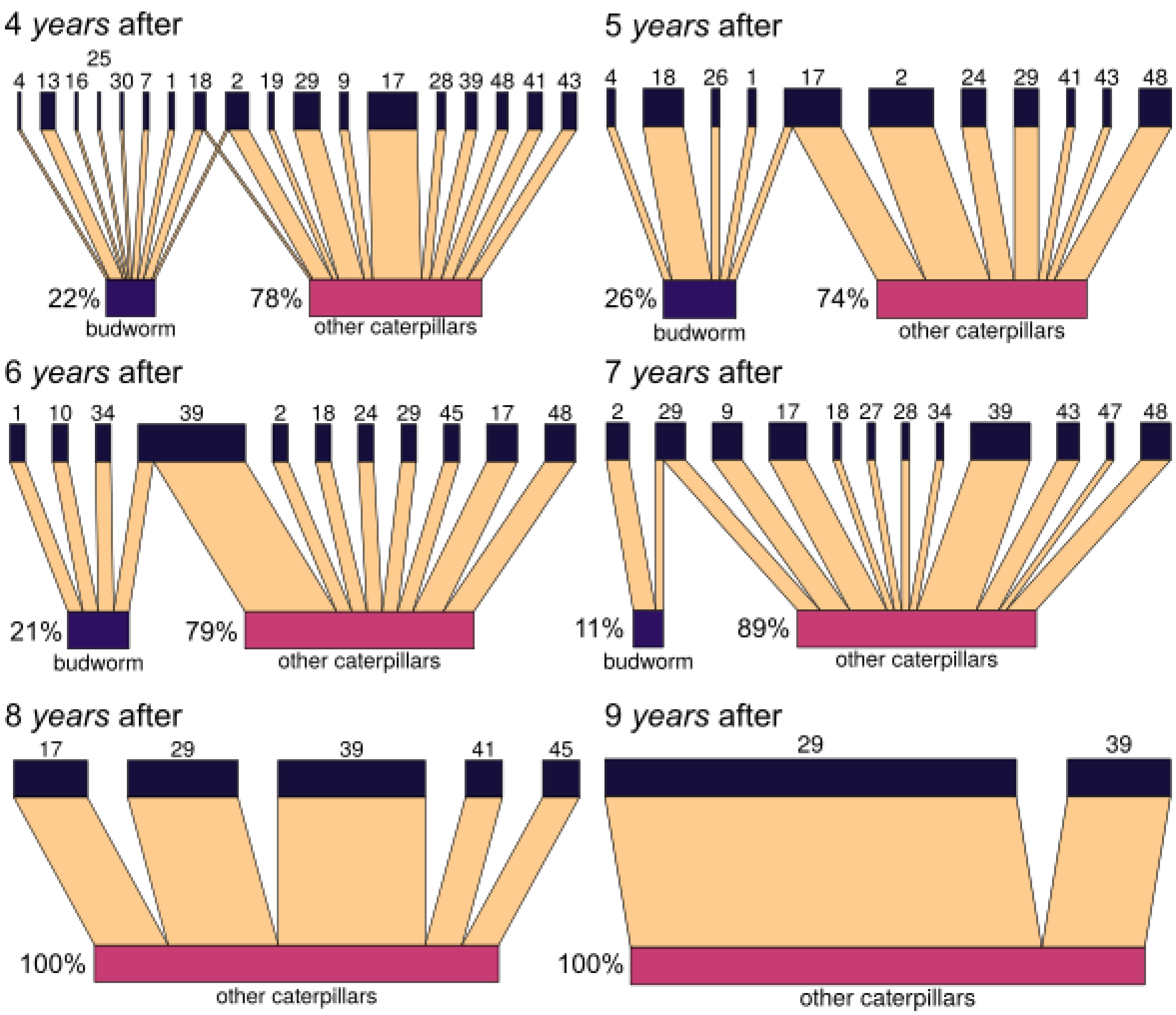


Figure S5 Graphical representations of the number of emergences of each parasitoid taxon (top boxes) from budworm and other caterpillar species (bottom boxes) over time. Six different relative years are shown: four relative years after the peak, five relative years after the peak, six relative years after the peak, seven relative years after the peak, eight relative years after the peak, and nine relative years after the peak.

Table S1 Parasitoid taxa found to attack budworm and other caterpillar species on balsam fir from this study compared to the corresponding parasitoid taxon found to attack budworm in Eveleigh et al. (2007)

| **Our parasitoids** | **Eveleigh et al. (2007) PNAS parasitoids** |
|---|---|
| 1. *Apanteles fumiferanae* | 9. *Apanteles fumiferanae* |
| 2. *Glypta fumiferanae* | 10. *Glypta fumiferanae* |
| 3. *Smidtia fumiferanae* | 1. *Smidtia fumiferanae* |
| 4. *Meteorus trachynotus* | 11. *Meteorus trachynotus* |
| 5. *Dirophanes hariolus* | 14. *Dirophanes hariolus* |
| 6. *Apechthis ontario* | 15. *Apechthis ontario* |
| 7. *Lypha fumipennis* | 2. *Lypha fumipennis* |
| 8. *Phyxe pecosensis* | 3. *Phyxe pecosensis* |
| 9. *Apanteles* sp. | 67. *Apanteles* sp. |
| 10. *Eumea caesar* | 4. *Eumea caesar* |
| 11. *Mesopolobus tortricus* | 13. *Mesopolobus tortricus* |
| 12. *Itoplectis conquisitor* | 16. *Itoplectis conquisitor* |
| 13. *Tranosema tenuifemur* | 66. *Synetaeris* sp. |
| 14. *Agria affinis* | 8. *Agria affinis* |
| 15. *Actia interrupta* | 6. *Actia interrupta* |
| 16. *Apanteles morrisi* | 19. *Apanteles morrisi* |
| 17. *Bassus binominata* | 20. *Bassus binominata* |
| 18. *Dolichogenidea absona* | 18. *Dolichogenidea absona* |
| 19. *Dirophanes* sp. | 48. *Phaeogenes* sp. |
| 20. *Madremyia saundersii* | 7. *Madremyia saundersii* |
| 21. *Scambus* sp. | 80. *Scambus* sp. |
| 22. *Macrocentrus linearis iridescens* | 41. *Macrocentrus linearis iridescens* |
| 23. *Enytus montanus* | 27. *Enytus montanus* |
| 24. *Charmon extensor* | 22. *Charmon extensor* |
| 25. *Apanteles petrovae* | 17. *Apanteles petrovae* |
| 26. *Tranosema rostrale* | 45. *Tranosema rostrale* |
| 27. *Chelonus* sp. | 29. *Chelonus* sp. |
| 28. *Diadegma pulicalvariae* | 50. *Diadegma pulicalvariae* |
| 29. *Orgilus* sp. | 21. *Orgilus* sp. |
| 30. *Apanteles milleri* | 38. *Apanteles milleri* |
| 31. *Sympiesis* sp. | 36. *Sympiesis* sp. |
| 32. *Stictophisthus flaviceps* | 93. *Stictophisthus* sp. |

| | |
|---|---|
| 33. *Pimpla pedalis* | 44. *Pimpla pedalis* |
| 34. *Meteorus versicolor* | 60. *Meteorus* sp. (versicolor?) |
| 35. *Elachertus cacoeciae* | 98. *Elachertus* |
| 36. *Aprostocetus esurus* | 35. *Aprostocetus* |
| 37. *Sarcophaga aldrichi* | 47. *Sarcophaga aldrichi* |
| 38. *Pristomerus* sp. | 30. *Pristomerus* sp. |
| 39. *Perilampus* sp. | 54. *Perilampus* sp. |
| 40. *Nemorilla pyste* | 5. *Nemorilla pyste* |
| 41. *Microgaster* sp. | 49. *Microgaster* sp. & 74. *Microgasterinae* |
| 42. *Lissonota acrobasidis* | 62. *Lissonota acrobasidis* |
| 43. *Glypta* sp. | 56. *Glypta* sp. |
| 44. *Exochus nigripalpis* | 37. *Bathythrix nigripalpis* |
| 45. *Diadegma* sp. | 39. *Diadegma* sp. |
| 46. *Ceromasia auricaudata* | Not in Eveleigh et al. (2007) PNAS parasitoids |
| 47. *Campoplex* sp. | 24. *Campoplex* sp. |
| 48. *Bassus dimidiator* | 40. *Bassus dimidiator* |
